# Heterogeneity of intrinsic excitability in Purkinje cells linked with longitudinal zebrin zones in the mouse cerebellum

**DOI:** 10.1101/2020.06.22.164830

**Authors:** Viet T. Nguyen-Minh, Khoa Tran-Anh, Izumi Sugihara

**Affiliations:** Department of Systems Neurophysiology, Graduate School of Medical and Dental Sciences, Tokyo Medical and Dental University, 113-8510, Tokyo, Japan; Center for Brain Integration Research, Tokyo Medical and Dental University, 113-8510, Tokyo, Japan

## Abstract

Heterogeneous populations of Purkinje cells (PCs), classified into zebrin-positive (Z+) and – negative (Z−) types, are arranged into separate longitudinal zones and are involved in different types of cerebellar learning. However, the electrophysiological phenotype that is brought about by the zebrin expression has not been much clarified in PCs. We compared electrophysiological characteristics in the soma and parallel fiber (PF)-PC synapse in Z+ and Z− PCs located in identified vermal and hemispheric zones in cerebellar slices in zebrin-reporter mice. Intrinsic excitability, intrinsic plasticity and PF-PC synaptic long term potentiation (LTP) occurred more strongly in Z− Purkinje cells than in Z+ PCs. The enhanced intrinsic plasticity was correlated with the reduction of medium-time-course after-hyperpolarization (mAHP) only in Z− PCs. These differences, which seem to be produced by the zebrin-linked expression of other functional molecules, may tune Z+ and Z− PCs to zone-specific cerebellar functions.

## Introduction

Neuronal mechanisms of long term information storage in the neuronal network include not only synaptic plasticity but also plasticity of the intrinsic excitability (intrinsic plasticity) of a neuron (Abraham, 2008; Titley et al., 2017). Indeed, intrinsic plasticity plays an essential role in the hippocampus for fear conditioning (McKay et al., 2009) and trace eyeblink conditioning (McEchron et al., 1997), in nucleus accumbens for cocaine addiction (Kourrich et al., 2015), and in the deep cerebellar nuclei for delay eyeblink conditioning (Wang et al., 2018). Neuronal intrinsic plasticity is often associated with a change in the amplitude of the afterhyperpolarization (AHP) which follows an action potential or other types of depolarization with various (fast, medium and slow) time courses. Voltage activated, calcium-activated potassium channels underlie the fast AHP and some of the medium AHP (Disterhoft and Oh, 2006).

Purkinje cells (PCs) are the sole output neuron in the cerebellar cortex. Purkinje cell activity can drive the induction of learned changes in behavior (Nguyen-Vu et al., 2013). Although PCs fire spontaneously, their firing frequency and pattern are dependent on and modulated by their excitability and various synaptic inputs (Zhou et al., 2014; Xiao et al., 2014; De Zeeuw and Brinke, 2015). Their inhibitory output then controls the firing pattern of target nuclear neurons (Person and Raman, 2011; White et al., 2014). Their excitability and various plasticity are deeply involved in cerebellar functions including control adaptation, learning motor performance and eyes movement (Boyden et al., 2004; Ito, 2013). Besides synaptic plasticities, the robust intrinsic plasticity of PCs, reported in in vivo and in vitro preparation (Belmeguenai et al., 2010; Grasselli et al., 2016; Shim, Jang et al; 2018), can have a significant effect through the PC output on the activity of cerebellar nuclear neurons (Wang et al., 2018; Person and Raman, 2011; White et al., 2014).

Noticeably, PCs are composed of heterogeneous populations that express several molecules at different levels (Cerminara et al., 2015; Hawkes, 2014). Since zebrin (zebrin II, aldolase C) is the representative of such molecules, PCs are generally classified into zebrin-positive (Z+) and zebrin-negative (Z−) populations (Brochu et al., 1990; Sugihara and Shinoda, 2004; Pijpers et al., 2006). Z+ and Z− PCs are distributed in different longitudinal zones to project to a particular subarea of the cerebellar nuclei and to be innervated by a particular subarea of the inferior olive according to the precise topographical connection (Sugihara and Shinoda, 2004; Pijpers et al., 2006; Oberdick and Sillitoe, 2011; De Zeeuw and Brinke, 2015). Thus, heterogeneous PCs generally belong to distinct functional networks (Sugihara and Shinoda, 2004; Aoki et al., 2019; Suzuki et al; 2012; Miterko et al., 2018). Indeed, spatial patterns of different types of climbing fiber responses in PCs are well linked with longitudinal or zebrin zones during behaving tasks with rewards in mice (Tsutsumi et al., 2015; Kostadinov et al., 2019; Tsutsumi et al., 2019).

Many other molecules including synaptic molecules are also expressed heterogeneously in the manner linked with zebrin expression (Hawkes, 2014; Beckinghausen and Sillitoe, 2019), suggesting the possibility that these molecules can affect electrophysiological properties of heterogeneous populations of PCs differently. However, the difference in electrophysiological properties including intrinsic plasticity has not been systematically studied between Z+ and Z− PC populations except that different firing frequencies of PCs in different areas have been reported in in vivo and in vitro preparations (Zhou et al., 2014; Xiao et al., 2014; Nguyen-Minh et al, 2019). Differences in synaptic properties have not been much explored between Z+ and Z− PCs either, beyond the report that the long term depression in the parallel fiber (PF)-PC synapse is more enhanced in anterior lobules which are rich in Z− PCs than in posterior lobules which are rich in Z+ PCs (Wadiche and Jahr, 2005).

Therefore, we aimed at examining intrinsic excitability, plasticity and related cellular physiological properties from Z+ and Z− PCs identified in neighboring zebrin zones. For this purpose, we used AldocV and other mice which allow clear PC type identification in vermal and hemispheric zones in slice preparation in which external inputs to cortical zones are eliminated.

## Results

### 1. Heterogeneous PCs in the slice preparation

AldocV mice have fluorescence expression in Z+ PCs (Fujita et al., 2014). Rgs8-EGFP mice seemed to have fluorescence expression in Z− PCs, which has not been much studied. Therefore, we examined the expression of aldolase C (zebrin) with immunostaining. The fluorescence expression pattern of Rgs8-EGFP mice was complementary with the zebrin expression pattern (Fig. 1A–D). Consequently, we could use both AldocV and Rgs8-EGFP mice to distinguish Z+ and Z− PCs in slices, avoiding artifacts due to particular genetic manipulation if any. To elucidate different characteristics between Z+ and Z− PCs, we focused on PCs in a pair of neighboring Z+ and Z− zones in one lobule to avoid possible factors related to lobular differences or vermis-hemisphere differences (Fig. 1E). In this study we used sagittal sections (Fig. 1F, J) and made recordings from PCs in zone 1+ and medial zone 1-, and lateral zone 1- and zone 2+, in vermal lobule IV-V in AldocV mice, and from PCs in zone 5+ and medial zone 5- in crus II in Rgs8-EGFP mice (Fig. 1F-M).

**Figure 1.**
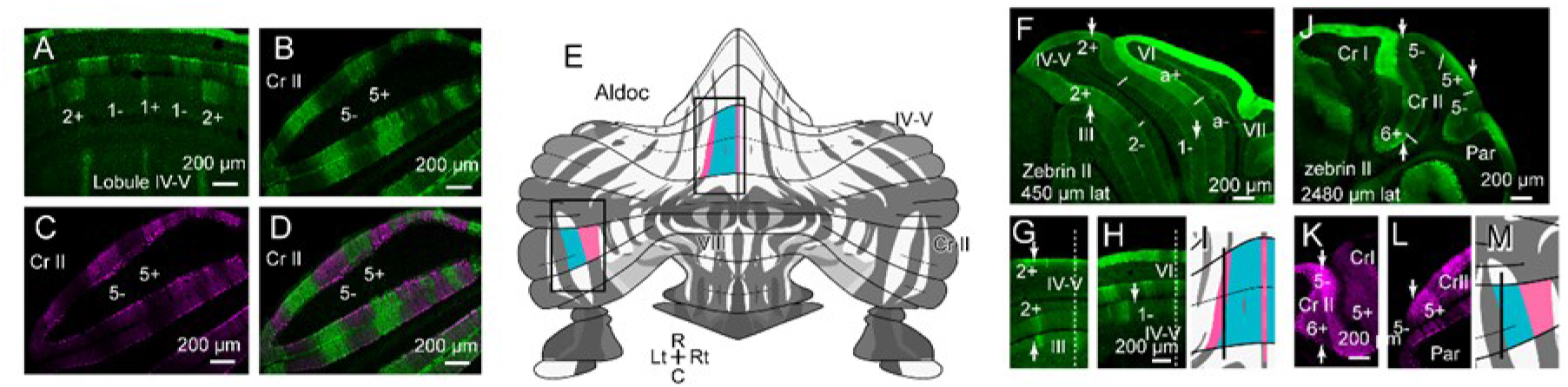
Identification of zebrin zones in AldocV mice and Rgs8-EGFP mice in coronal and sagittal sections. (A-D) coronal sections of vermal lobules IV-V and VIa (A) in the AldocV mouse and at left crus II (B-D) in the Rgs8-EGFP mouse at postnatal day 20 (P20). Immunostaining for zebrin II (alsolase C), EGFP fluorescence, and merged image are shown for the crus II section (as B, C and D, respectively). (E) Recording areas are mapped in the scheme of zebrin zones in the cerebellar cortex (Sarpong et al., 2018). (F) Sagittal section 450 μm lateral to the midline showing zebrin zones in lobules IV-V and neighboring lobules in the aldoc V mouse. Short white lines indicate approximate boundaries of zebrin zones. Paired (downward and upward) arrowheads and single downward arrowhead indicate the positions of panels (G) and (H), respectively. (G, H) Coronal section at the levels indicated by arrows in (F). White dashed lines indicate position of the midline. Arrowheads indicate the position of panel (F). (I) Identified zebrin zones in (F) shown by colors in the unfolded scheme. (J-M) Images of crus II and neighboring lobules 2480 μm lateral to the midline, arranged in a similar way to (F-I). Abbreviations, IV-VIII, lobule names; 1+, 1-, a+, a-, 2+, 2-, 5+, 5-, zebrin zones 1+ and so on; C, caudal; Cr I, crus I; Cr II, crus II; Lt, left, Par, paramedian lobule; Rt, right, R, rostral.

### 2. Intrinsic excitability is enhanced in the Z− zones

To examine the intrinsic excitability of PCs, the number of spikes evoked by 500 ms current injection was plotted against the intensity of injected current (spike-current relationship) for Z+ and Z− PCs in the three areas (Fig. 2A–C). The spike count showed a similar increase in Z+ and Z− PCs for small current injections (0–200 pA), whereas it was higher in Z− PCs than in Z+ PCs for stronger current injections (300–500 pA) (p<0.001, p<0.01, p<0.001, two-way ANOVA with repeated measurement 0–500 pA, n=18, 18, 13, 13, 13, and 13 PCs in zones 1+, medial zone 1-, lateral zone 1- and zone 2+ in lobule IV-V, and zones 5+ and 5- in crus II, respectively).

**Figure 2.**
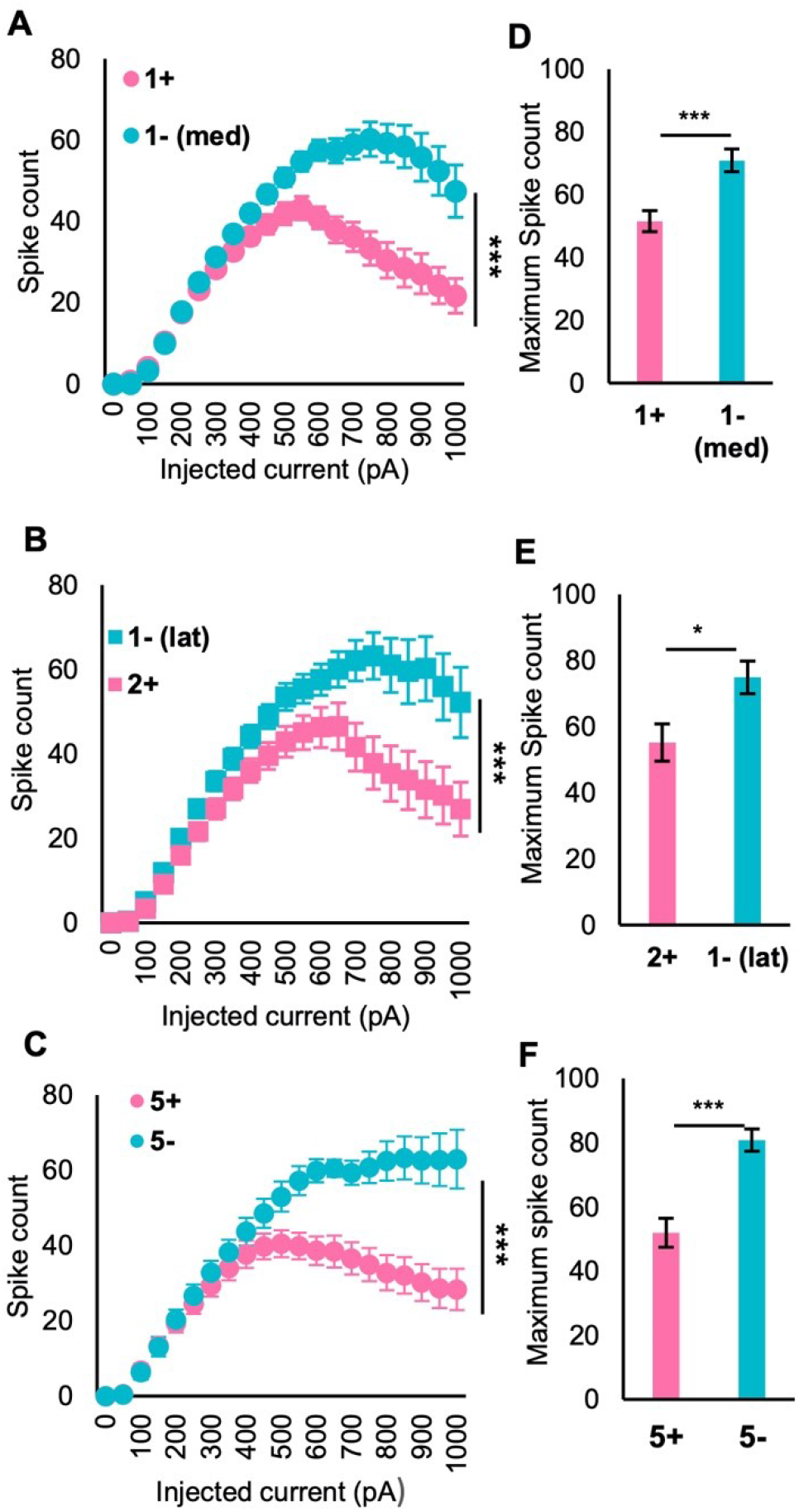
Comparison of the intrinsic excitability between Z+ and Z− PCs in neighboring zones in vermal and hemispheric lobules. (A-C) Plot of spike-current relationships, i.e., the spike count measured during 500 ms square current injection of various intensity under current clamp mode in 18, 18 (A), 13, 13 (B), 14 and 14 (C) PCs. (A, zebrin zones 1+ and lateral 1- in lobule IV-V; B, lateral 1- and 2+ in lobule IV-V; C, 5+ and 5- in crus II). (D-F) Comparison of the maximum spikes count between Z+ and Z− PCs in the regions shown in A, B, C respectively. AldocV mice were used in A and B, and Rgs8-EGFP mice were used in C. Mean ± SEM are shown in all panels. ***p<0.001, Two-way ANOVA with repeated measures (n=18+18, 13+13, 14+14 in A, B, C, respectively), Student t-test (n=18+18, 13+13, 14+14 in D, E, F, respectively).

A larger current injection (500–800 pA) achieved spike count saturation as reported in mouse PCs (Yamamoto et al., 2019). Z+ PCs showed saturation around a frequency of 40 per 500ms with 500-600 pA, while Z− PCs showed saturation around 60 per 500 ms with 700–800 pA, in all the three areas (Fig. 2A–C) (p<0.001, two-way ANOVA with repeated measurement 0-1000pA, n=18, 18, 13, 13, 13, and 13 PCs in zones 1+, medial zone 1-, lateral zone 1- and zone 2+ in lobule IV-V, and in zones 5+ and 5- in crus II, respectively). The maximum spike count was about 1.5 times higher in Z− PCs than in Z+ PCs in all areas. Specifically, the maximum spike count was 70.9 ± 3.6, 75 ± 4.9, 80.9 ± 3.5 in Z− zones (medial 1-, lateral 1- and 5-), whereas the maximum spike count were 51.6 ± 3.4, 55.2 ± 5.6, 52.0 ± 4.5 in Z+ zones (1+, 2+ and 5+) (Fig. 2D–F). Thus, Z− PCs were more excitable and able to fire more frequently than Z+ PCs. We did not observe significant differences in the input resistance or capacitance between Z+ and Z− PCs in the present study as well as in our previous study in lobule VIII (Nguyen-Minh et al., 2019).

Inactivation of repetitive firing at strong current injection, as indicated by the cease of the firing during 500 ms current injection, was more prominent in Z+ PCs than in Z− PCs in all the three areas (cf. Fig. 4B, C top traces). The intensity of current injection that brought about the maximum number of spikes was larger in Z− PCs than in Z+ PCs also in the three areas (data not shown).

These results indicated higher excitability of Z− PCs than in Z+ PCs. It was in line with similar findings obtained in the different cerebellar areas (Nguyen-Minh et al., 2019) and also with the in vivo finding of general higher simple spike firing rate in Z− PCs than in Z+ PCs in various areas of the rat and mice cerebellum (Zhou et al., 2014; Xiao et al., 2014; Wu et al., 2019). Furthermore, the results indicated that Z− PCs were capable of translating responsible to for stronger current injections than Z+ PCs.

### 3. Subthreshold AHP correlated with excitability in Z− PCs but not in Z+ PCs

The higher capability of repetitive spike firing in Z− PCs than in Z+ PCs indicates some difference in depolarization-evoked ionic conductances between these PCs. The depolarization-evoked ionic conductances are reflected in the AHP. The intensity of the AHP can be measured more simply and accurately in the AHP evoked by the subthreshold depolarization (“subthreshold AHP”) than the AHP evoked by simple or complex spikes in PCs or post-burst AHP (Belmeguenai et al., 2010; Grasselli et al., 2016; Paukert et al., 2010). We, therefore, compared subthreshold AHP to elucidate mechanisms for different repetitive spike firing responses between Z+ and Z− PCs.

We analyzed differences in the subthreshold AHP evoked by synaptic and current-injection protocols. In the synaptic protocol, PFs were stimulated with a train of 5 stimuli. AHP/EPSP ratio, which represented the size of the subthreshold AHP, was significantly higher in Z+ PCs than in Z− PCs in both pairs of neighboring Z+ and Z− zones measured in lobule IV-V (Fig. 3A; 18.2 ± 1.4 %, n=18 in 1+ vs. 12.9 ± 0.7 %, n=18 in medial 1-, p<0.01, unpaired Student’s t-test; 17.2 ± 1.3%, n=13 in 2+ vs. 13.0 ± 1.1 %, n=13 in lateral 1-, p<0.05, unpaired Student’s t-test, Fig. 3C). Data in two pairs of Z+ and Z− zones in lobule IV-V distributed in the same range and combining them also indicated a significant difference in the AHP/EPSP ratio (p<0.001, unpaired Student’s t-test, n=31,31). To examine the contribution of the subthreshold AHP to the repetitive firing capability of PCs, the maximum spikes count (Fig. 2D–F) was plotted against the AHP/EPSP ratio for Z+ and Z− PCs (Fig. 3E, G). Subthreshold AHP correlated negatively with the maximum spike count in Z− PCs (Pearson’s r = −0.53, p =0.002, Fig. 3G). On the contrary, there was no correlation in Z+ PCs (Pearson’s r = −0.09, p = 0.62, Fig. 3E). The results suggested that subthreshold AHP correlated with the intrinsic firing only in Z− zones.

**Figure 3.**
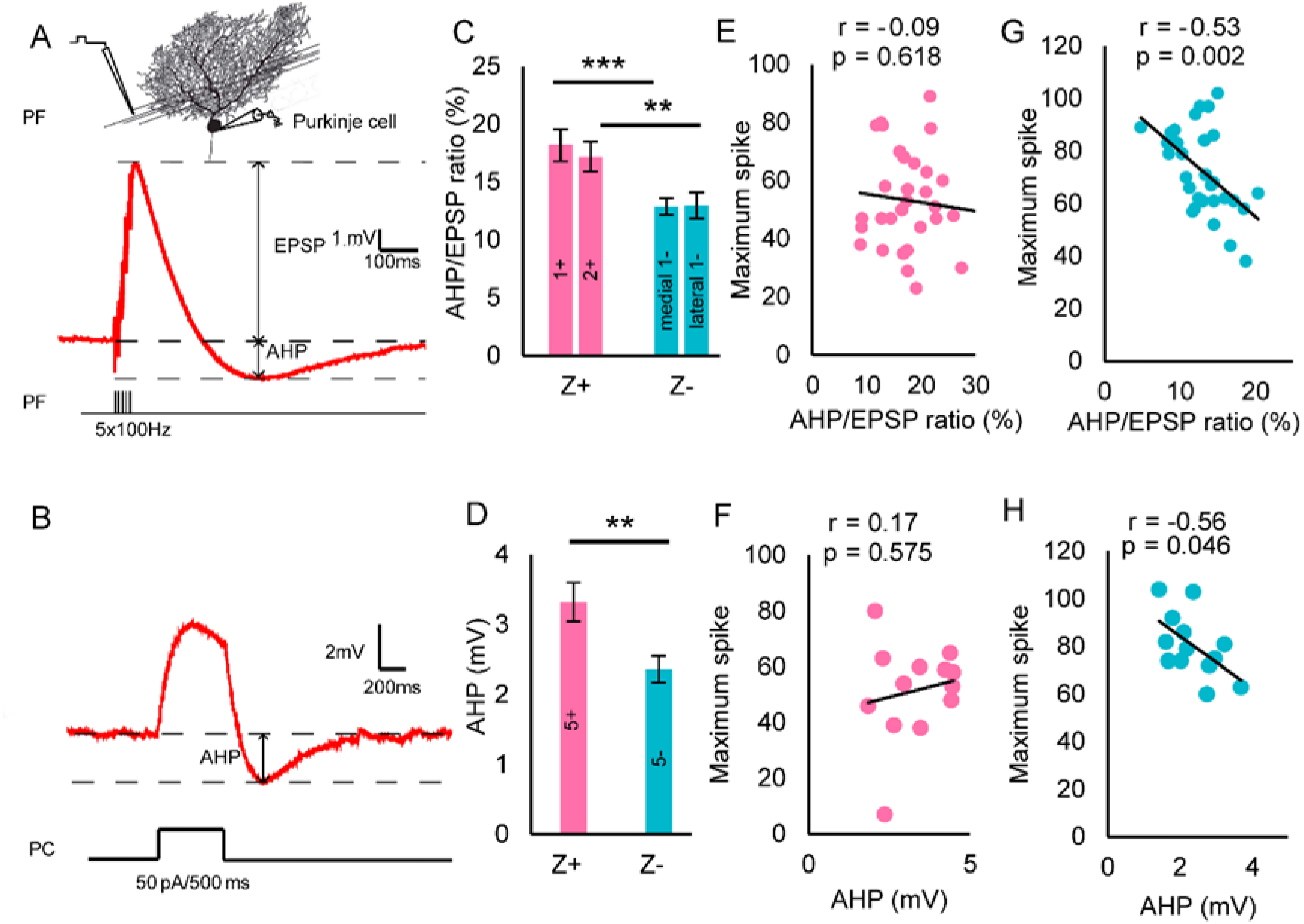
Comparison of the afterhyperpolarization (AHP) intensity between Z+ and Z− PCs in neighboring zones in vermal and hemispheric lobules. (A, B) Recordings of membrane potential to illustrate measurements of the AHP intensity. The synaptic protocol, consisting of five repetitive PF stimuli at 100 Hz (A), was used in PCs in zebrin zones 1+, medial 1-, lateral 1- and 2+ in lobule IV-V in AldocV mice. The current injection protocol, consisting of a square current injection of 50 pA for 500 ms (B), was used in PCs in zebrin zones 5+ and 5- in crus II in Rgs8-EGFP mice. (C) Comparison of the AHP intensity in lobule IV-V. AHP intensity was represented by the ratio (in percent) of the AHP peak amplitude to the EPSP peak amplitude obtained with the synaptic protocol. (D) Comparison of the AHP intensity in crus II. AHP intensity was represented by the AHP amplitude obtained with the current injection protocol. ***p < 0.001; **p < 0.01. Mean ± SEM are shown in C and D. (E–H) The relationship between the AHP intensity and maximum spike count in Z+ and Z− PCs in lobule IV-V (E and G, in AldocV mice), and in crus II (F and H, in RGs8-EGFP mice). Regression analysis was performed to get Person’s r value and p value for the data in each panel.

We then made a similar analysis about the subthreshold AHP with the current injection protocol (Schreurs et al., 1998) in PCs in crus II (Fig. 3D). The AHP amplitude of Z− PCs (2.16 ± 0.19 mV) was significantly smaller than that of Z+ PCs (3.32 ± 0.27 mV) (p<0.01, Student’s t-test, n=13 each group) (Fig. 3D). The plots of the maximum spike count against the subthreshold AHP amplitude in individual PCs showed negative correlation in Z− PCs (Pearson’s r = −0.56, p =0.046, Fig. 3H), whereas the same plot showed no correlation in Z+ PCs (Pearson’s r = – 0.17, p =0.58, Fig. 3F).

A smaller AHP was observed in Z− PCs than in Z+ PCs in both protocols were likely to reflect higher intrinsic excitability of Z− PCs (Fig. 2A-C). Furthermore, the results indicated that the variation of AHP intensity correlated more with the variation of intrinsic excitability in individual Z− PCs than in individual Z+ PCs.

### 4. LTP of intrinsic excitability more enhanced in Z− zones than in Z+ zones

The intrinsic excitability of PCs, represented by the increase of spike firing frequency in response to the increase of injected current, is potentiated permanently by tetanizing stimulation (long term potentiation of intrinsic excitability or LTP-IE; Belmeguenai et al., 2010). However, this property has not been compared between Z+ and Z− PCs. The tetanizing stimulation (“LTP-IE protocol”, Fig. 4A, Shim, Jang et al; 2018) enhanced intrinsic excitability in both Z+ and Z− PCs in vermal lobule IV-V (Fig. 4B, C, F, G). To make a detailed comparison between Z+ and Z− PCs, we measured the current-spike relationship before (“baseline”) and 10-20 min after the LTP-IE protocol (“10 min”, and “20 min”, Fig. 4F, G). In control experiments, in which no LTP-IE protocol was given, the current-spike relationship was stable among “baseline”, “10 min” and “20 min” in both Z+ and Z− PCs (Fig. 4D, E). However, applying the LTP-IE protocol induced enhancement in the current-spike relationship and this enhancement remained for 20 min (Fig. 4F, G, top). This enhancement was statistically significant in both PCs in Z+ (Fig. 4F top, p<0.001, two-way ANOVA with repeated measure, n=13) and Z− zones (Fig. 4G top, p<0.001, two-way ANOVA with repeated measure, n=13). Noticeably, the maximum spike count was also increased.

**Figure 4.**
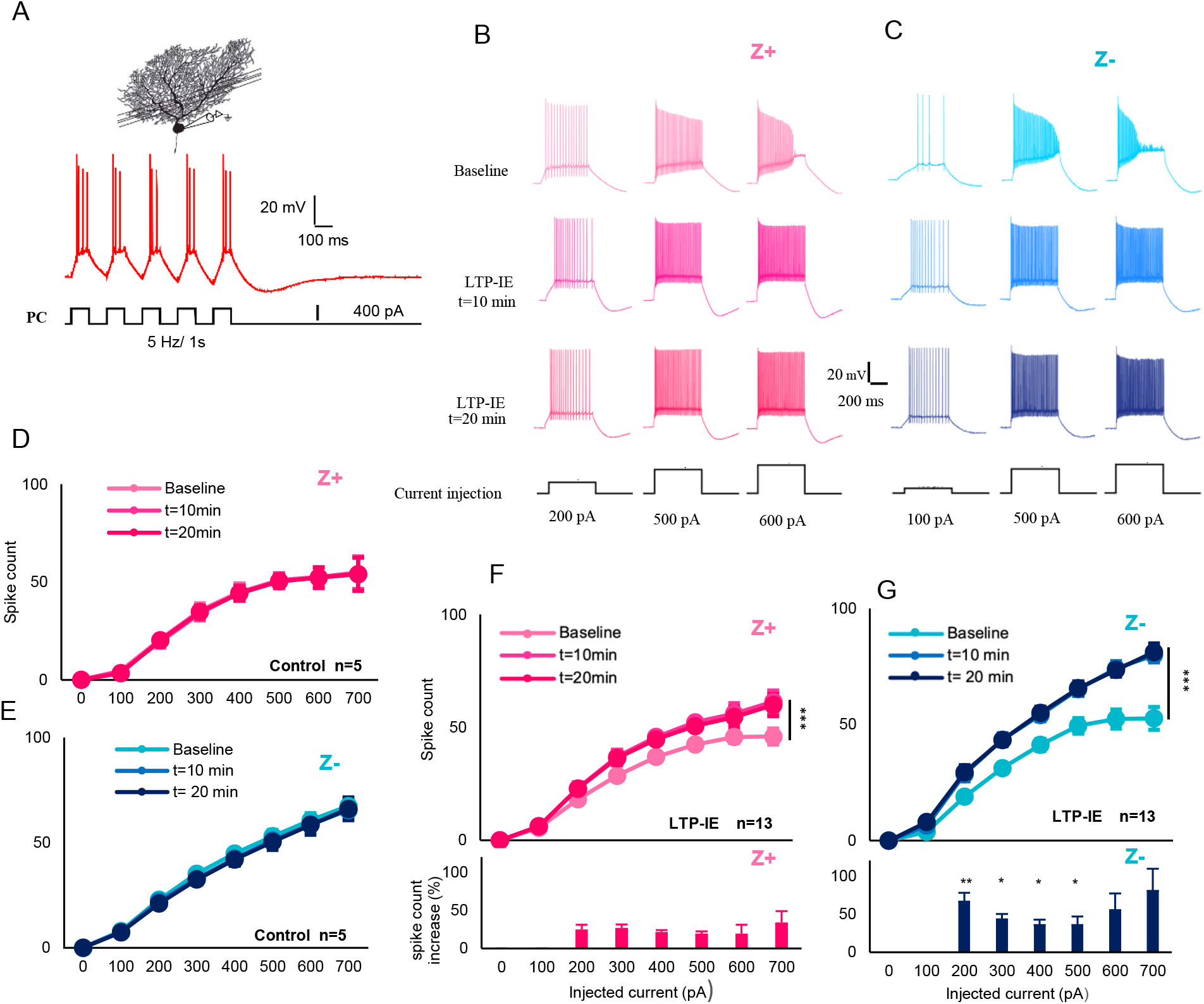
Comparison of the intrinsic plasticity between Z+ and Z− PCs in neighboring zones. (A) The direct stimulation protocol to induce the LTP-IE or intrinsic plasticity. A run of five square-shaped current injection (400 pA, 100 ms ON and OFF, 5 Hz, as shown in the panel) was repeated every 2 s for 5 min. (B, C) Spike response to square current injection (100, 500 and 600 pA for 500 ms) before, and 10 and 20 min after giving the LTP-IE protocol in PCs in zone 1+ (B) and medial zone 1- (C) in lobule IV-V respectively. (D–G) Potentiation of the current-spike relationship in Z+ (D, F) and Z− (E, G) PCs. The number of spikes during 500 ms square current injection of various intensity (100–700 pA) was plotted against the intensity of injected current. In the control experiment without LTP-IE (n=5 PCs in D, 5 PCs in E), initial measurement and two measurements 10 and 20 minutes after the first measurement were superimposed. In the LTP-IE experiments (n=13 PCs in F, 13 PCs in G), initial measurement and two measurements 10 and 20 minutes after the LTP-IE protocol (tetanization) were superimposed. Increase of the spike count (F bottom, G bottom) was then calculated from the measurements before and 20 minutes after the LTP-IE protocol. Mean ± SEM are shown. In (F top), (G top) change of the spike number was tested by two-way ANOVA with repeated measures between baseline and t=20 min. In the F, G bottom panel, the difference in the spike count increase between Z+ (F) and Z− (G) PCs was tested with Mann-Whitney U test. ***p < 0.001; **p < 0.01; *p < 0.05.

Next, we compared the degree of the LTP-IE between Z+ and Z− PCs. The increase of the spike count, which was measured in percent at 20 min after the LTP-IE protocol from the baseline spike count with various intensity of current injection, was 1.5~3 fold stronger in Z− PCs than in Z+ PCs at 200, 300, 400, 500 pA current injection (Fig 4F, G bottom, p=0.0012, p=0.012, p=0.02, p=0.026, respectively, Mann-Whitney U test, n=13). The result indicated a higher degree of LTP-IE in Z− PCs than that in Z+ PCs.

### 5. Distinct modulation of different AHP phases underlying the LTP-IE in Z+ and Z− PCs

Intrinsic plasticity is brought about by the AHP depression in PCs (Belmeguenai et al., 2010). Since different phases of AHP are separately modulated and involved in changing the intrinsic excitability (Disterhoft and Oh, 2006), we next investigated the relationship between intrinsic plasticity and changes in different phases of AHP in Z+ and Z− PCs. We classified the AHP into three types, i.e. fast, medium and slow AHPs (fAHP, mAFP, sAFP), according to Disterhoft and Oh (2006). While the fAHP occurred after a single spike with a latency of less than a few milliseconds, the mAHP and sAHP occurred after a burst of spikes with a latency of 100–200 ms and 600–700 ms (Fig 5A, B). These different phases of AHP reflect different underlying mechanisms (ionic channels and ionic pumps, Disterhoft and Oh, 2006) and contribute to intrinsic plasticity differently (Belmeguenai et al., 2010).

**Figure 5:**
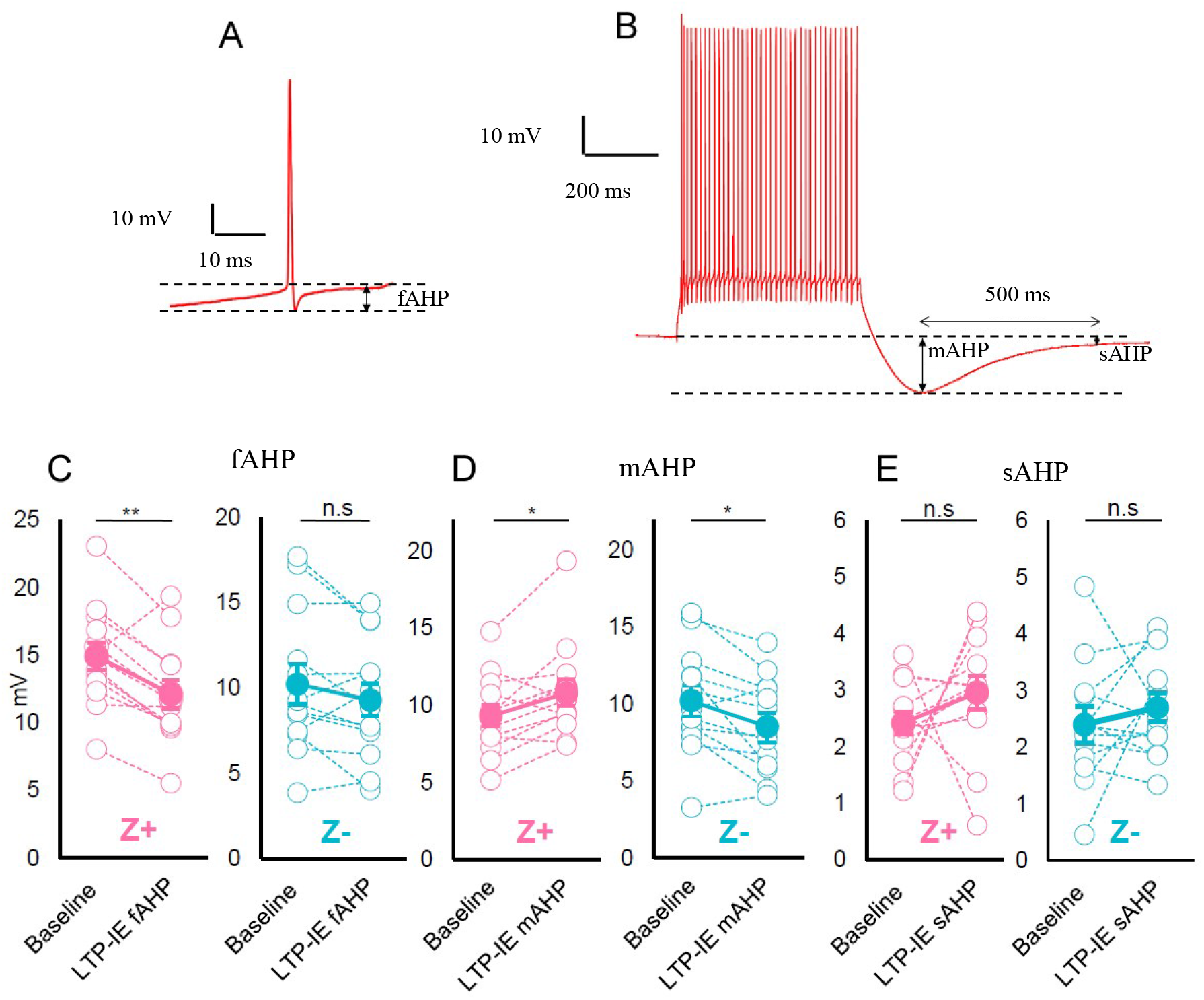
Distinct modulation of different AHP phases underlying the LTP-IE in Z+ and Z−PCs. (A) Sample trace to show measurement of the the fast AHP (fAHP) (B, C) Sample trace to show measurement of the middle and slow AHP (mAHP and sAHP). (C–E) Change of three phases of the AHP after the LTP-IE protocol (tetanization). Amplitudes of the fAHP (C), mAHP (D) and sAHP (E) were compared before and 20 minutes after the LTP-IE protocol in 13 Z+ (left graph in each panel) and 13 Z− (right graph in each panel) PCs. (p=0.001, 0.1, 0.015, 0.013, 0.26, 0.39 in C left, C right, D left, D right, E left and E right, respectively, Student’s paired t-test, n=13 in all) ***p < 0.001; **p<0.01; *p < 0.05.

We compared amplitudes of fAHP, mAHP and sAHP between the baseline period before giving the LTP-IE protocol (tetanizing stimulation) and 20 minutes after the LTP-IE protocol in Z+ and Z− PCs in vermal lobule IV-V (Fig. 5C–E). Z+ PCs showed decrease in fAHP (Fig. 5C, Left, p<0.001, paired Student’s t-test), increase in mAHP (Fig. 5D, Left, p<0.05, paired Student’s t-test) and no significant change in sAHP (Fig. 5E, Left, p>0.05, paired Student’s t-test). However, Z− zones PCs showed a decrease in mAHP (Fig. 5D, Right, p<0.05, paired Student’s t-test) and no significant change in fAHP (Fig. 5C, Right, p>0.05, paired Student’s t-test) or in sAHP (Fig. 5E, Right, p>0.05, paired Student’s t-test). Our results indicated that LTP-IE protocol (tetanizing stimulation) modulate distinct phases of AHP differently in Z+ and Z− PCs. The decrease in mAHP and the decrease in fAHP were likely to be responsive to the intrinsic plasticity in Z+ and Z− PCs, respectively.

### 6. Different degree of PF-PC synaptic LTP between Z+ and Z− PCs

The Intrinsic plasticity shares the same molecular pathway as the postsynaptic pathway of LTP of the PF-PC synaptic transmission in PCs (Belmeguenai et al., 2010). The different efficiency of intrinsic plasticity between Z+ and Z− PCs (above) suggested the possibility of different efficiency in the synaptic LTP between these PCs. Since we primarily focused on postsynaptic factors in Z+ and Z− PCs, we used the stimulation protocol that minimizes the effects of presynaptic factors, which may also affect differences between Z+ and Z− PCs. First, we used parasagittal slices, rather than transverse slices, to bypass the involvement of the presynaptic NMDA receptor in PFs (Bouvier et al., 2016). Secondly, we applied low-frequency stimulation of PFs to reduce the nitrogen monoxide synthesis in molecular layer interneurons (Wang et al., 2014).

In the middle of continuous monitoring of PF-PC synaptic excitatory postsynaptic current (EPSC), PF stimulation frequency was increased to 1 Hz for 5 min to induce synaptic LTP in Z+ and Z− PCs in zones 1+, 1- and 2+ in lobule IV-V (Fig. 6A, B). The potentiation percentage, i.e. the ratio of the average EPSC 20-25 min after the PF stimulation to the average EPSC before the PF stimulation, was significantly higher in Z− PCs than in Z+ PCs (Fig. 6C, p<0.05, Student’s t-test, n=6). The results demonstrated that LTP in the PF-PC synaptic transmission was more enhanced in Z− PCs than in Z+ PCs.

**Figure 6.**
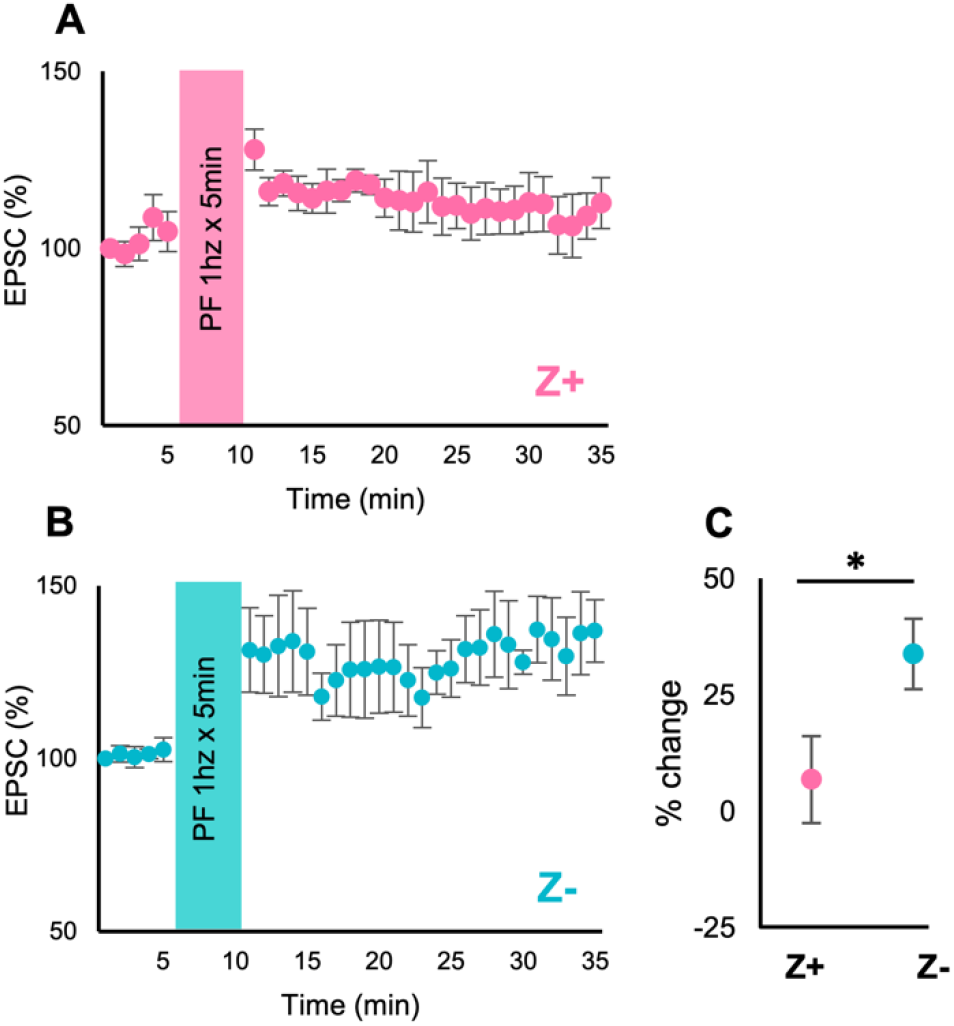
Comparison of the LTP in the PF-PC synaptic transmission between Z+ and Z− PCs in neighboring zones. (A, B) Time graph of the normalized EPSC amplitude before and after the 1-Hz PF stimulation in six Z+ PCs in zebrin zone 1+ and 2+ (A) and in six Z− PCs in zebrin zone 1- in lobule IV-V(B). (C) Comparison of the EPSC amplitude 21–25 min after the 1-Hz PF stimulation. *p<0.05 Student t-test. Mean ± SEM were plotted in all panels.

## Discussion

The present study carefully compared intrinsic excitability and PF-PC synaptic properties between Z+ and Z− PCs in neighboring zebrin zones in the slice preparation. Z− PCs were more enhanced in intrinsic excitability, intrinsic plasticity, and PF-PC synaptic LTP. The intrinsic plasticity was mediated by different mechanisms in Z− and Z+ PCs. As a whole, the present study demonstrated that cerebellar PCs are heterogeneous not only in molecular expression profiles and topographic axonal connection patterns but also in basic electrophysiological properties as well as in the synaptic plasticity.

### Electrophysiological heterogeneity in PC populations

Since the discovery of zonal distribution of the heterogeneous population of PCs which differs in expression levels of several molecules, it has been a long-lasting question of what different properties such PC populations have at the cellular physiological level. By sampling Z+ and Z− PCs from identified neighboring zones in an identified lobule in the slice preparation from mice with fluorescent labeling of specific populations of PCs, we could eliminate possible effects of environmental factors and different afferent inputs. This is the ideal situation to focus on different cellular physiological properties between Z+ and Z− PCs. Such a situation was not necessarily attained in previous in-vivo and in-vitro studies about heterogeneous physiological properties in PCs (Wu et al., 2019; Wadiche and Jahr, 2005). Our previous study with similar slice preparation from AldocV mice has shown no difference in input resistance or capacitance or basic properties in PF-PC synaptic transmission between Z+ and Z− PCs but indicated evidence of different intrinsic excitability between these PCs (Nguyen-Minh et al, 2019), which has led us to the present study.

PCs are capable of the repetitive firing of simple spikes. Sampling PCs in various cerebellar areas in in vivo preparation showed that Z− PCs generally shows a higher frequency of simple spike firing (Zhou et al., 2014; Xiao et al., 2014). Although these studies could not exclude the possibility of different intensities of excitatory inputs between these PCs, the present study confirmed that Z− PCs have higher intrinsic excitability and a smaller AHP, which underlie the higher intrinsic excitability, than Z+ PCs. Underlying mechanisms were not clarified yet, the difference in some voltage-dependent ionic currents was likely to be involved, since, the reduction of K^+^ conductance increase neuronal excitability by decreasing AHP (Abraham, 2008; Disterhoft and Oh, 2006; Belmeguenai et al., 2010).

Furthermore, Z− PCs showed higher intrinsic plasticity; a tetanization increased the intrinsic excitability more in Z− PCs than in Z+ PCs. Underlying mechanisms for the intrinsic plasticity were different between Z− and Z+ PCs; a decrease in mAHP occurred in Z− PCs whereas a decrease in the fAHP occurred in Z+ PCs. Besides intrinsic plasticity, the LTP of the PF-PC synaptic transmission was also stronger in Z− PCs than in Z+ PCs. As a whole, Z− PCs are more sensitive to tonic excitatory synaptic input and respond to a higher frequency of firing and more sensitive to the enhancement of simple spike activity than Z+ PCs.

### Functional significance of the electrophysiological heterogeneity in cerebellar zones

Z− and Z+ PCs are separately distributed in tens of longitudinal zones in the cerebellar cortex. Each zone has a topographic connection with particular subareas of the inferior olive through climbing fibers and with particular subareas of the cerebellar nucleus through PC axons (Sugihara and Shinoda, 2004; Pijpers et al., 2006). Zones of Z+ PCs (or Z+ zones) tend to receive climbing fiber input from regions of the inferior olive that are dominated by descending inputs, whereas Z− zones receive climbing fiber input from regions of the inferior olive that receive predominantly peripheral inputs (Cerminara et al., 2015; Sugihara and Shinoda, 2004; Voogd and Ruigrok., 2004). Indeed, Ca^2+^ imaging study reported CF-dependent activity at different phases in Z+ and Z− zones in crus II, in which both Z+ and Z− zones occupy comparable widths, in behaving mice (Tsutsumi et al., 2015; Tsutsumi et al., 2019). Cortical areas that control somatosensory reflexes such as eye blinking reflex (hemispheric lobule V and simple lobule) and locomotion (vermal lobules I-VIa) are mostly composed of wide Z− zones and narrow Z+ zones, while cortical areas that control eye movements (flocculus and nodulus), posture, and cognitive function (crus I and vermal lobules VI-VII) are mostly composed of wide Z+ zones. Thus, different types of PC excitability and plasticity may be well-tuned for different types of learning and adaptation mediated in these areas. Indeed, one such idea has been proposed in adaptation mechanisms of eyeblink response and vestibule-ocular response (De Zeeuw and Brinke, 2015). The quick and robust plasticity in Z− PC supports the “temporal coding” mechanism (the spikes occur at millisecond precision) of eye-blink conditioning between unrelated unconditioned and conditioning stimuli, whereas the small plasticity in Z+ may fit with “rate coding” in the vestibulo-ocular reflex adaptation that requires precise adjustment of correlated movements between the head and eye (De Zeeuw and Brinke, 2015; Payne et al., 2019). The present results propose that the more enhanced intrinsic plasticity and synaptic LTP in Z− zones than in Z+ zones, as revealed in the present study, may also be considered in understanding area-dependent functional differences of the cerebellum.

### Molecular mechanisms underlying the electrophysiological heterogeneity

Many molecules involved in controlling neuronal excitability and synaptic transmission have heterogeneous expression profiles linked to the zebrin expression patterns (Cerminara et al., 2015; Hawkes, 2014); Z+ PCs preferentially express PKCδ, EAAT4, and PLCβ3, whereas Z− PCs preferentially express PLCβ4 and TRPC3 (only in the vermis, Wu et al., 2019). These differences in molecular expression suggest different electrophysiological properties between Z+ and Z− PCs (Fig. 7). In the tetanic PF stimulation for the LTP of PF-PC synaptic transmission, the lack of EAAT4 in Z− PCs induces higher activation of mGluR1 in the extrasynaptic area in Z− PCs (Wadiche and Jahr, 2005). The mGluR1 activation, mediated by Gq, induces PLCβ4 activation, which then induces inositol-1,4,5-triphosphate (IP3) -dependent Ca^2+^ release from the intracellular Ca^2+^ storage. Interaction of the mGluR1b receptor with the Ca^2+^-permeable TRPC3 channel (Wu et al., 2019) is another possible pathway for Ca^2+^ increase in Z− PCs. The LTP-IE protocol can also induce a larger increase of intracellular Ca^2+^ concentration in Z− PCs because of larger influx through voltage-activated Ca^2+^ channels due to more frequent action potentials in Z− PCs. Ca^2+^ then activates PLCβ4 further and also the protein phosphatase 1 (PP1), protein phosphatase 2A (PP2A) and protein phosphatase 2B (PP2B), which induce LTP of the PF-PC synaptic transmission (Belmeguenai et al., 2010), and also LTP-IE by suppressing the small conductance Ca^2+^-activated K channel (SK channel) putatively involved in the mAHP (Belmeguenai et al., 2010; Grasselli et al., 2020; Titley et al., 2020). Thus, intrinsic excitability and synaptic LTP can be more enhanced in Z− PCs. On the contrary, in Z+ PC, PLCβ3-induced diacylglycerol (DAG) release can activate PKCδ, which requires DAG but not Ca^2+^ for activation (Rosse, et al., 2010). PKCδ might modulate the large-conductance Ca^2+^-activated K channels (BK channel), contributing to the synaptic LTP and LTP-IE (Helene et al., 2003; Barmack et al., 2001) to some extent. Moreover, PKC phosphorylates Ser1506 of *α* subunit of the voltage-gated Na^+^ channel to slow its inactivation (Li et al., 1993), which might support the continueous firing of PCs under large current injections. Thus, we speculate that the reported differences in the molecular expression profiles are correlated with the differences in electrophysiological properties that we observed in the present study between Z− and Z+ PCs.

**Figure 7.**
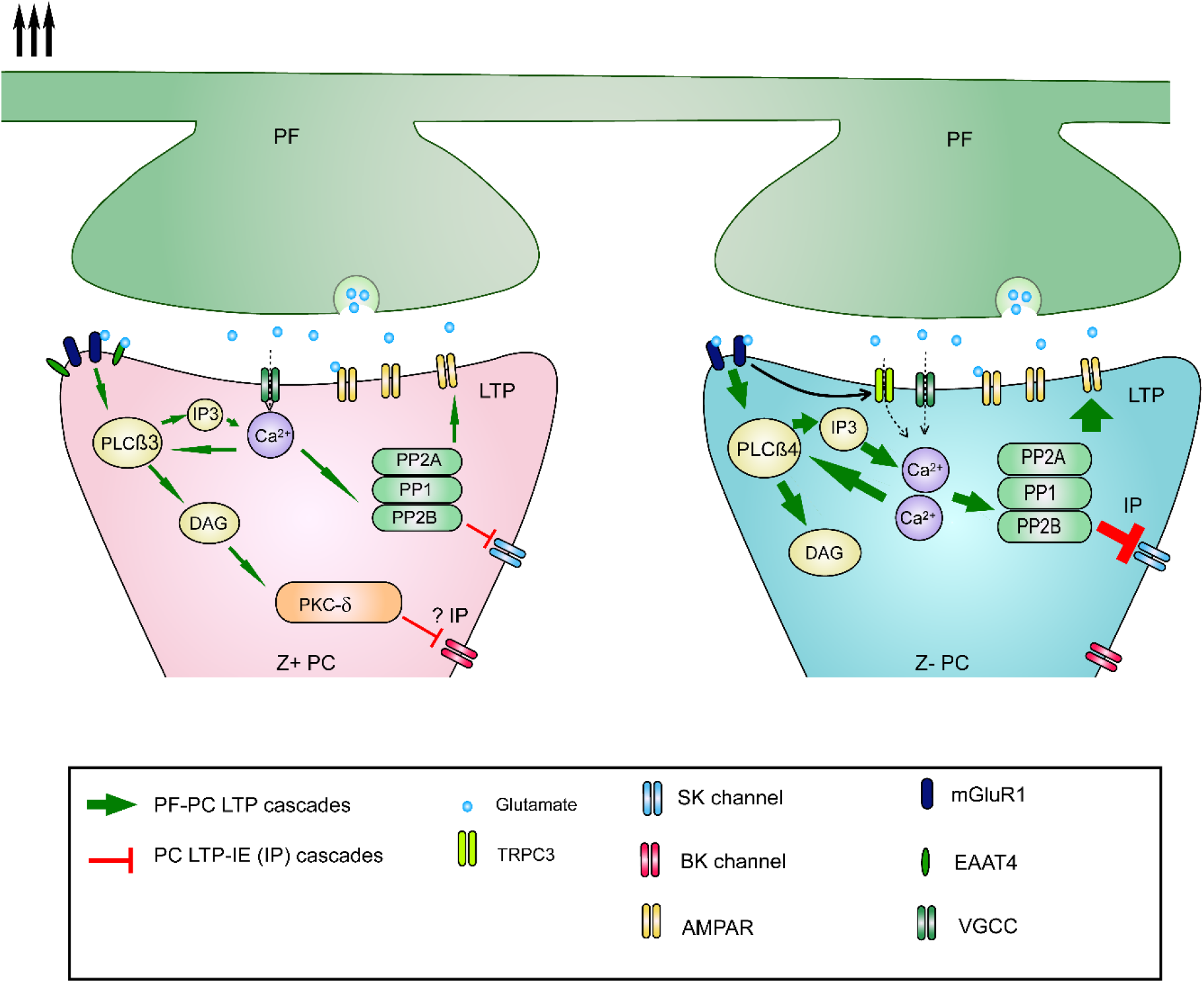
Putative molecular mechanisms underlying synaptic and intrinsic plasticity in Z+ and Z− PCs. Pathways involved in LTP at the PF-PC synapse are colored in green, while pathways involved in Intrinsic plasticity, which are not necessarily localized in the spine, are colored in red. Refer to the last section of Discussion for an explanation Abbreviations: PF, Parallel Fibers; PC, Purkinje cells; AMPAR, AMPA receptor; PKC, protein kinase C; PLC, phospholipase C; PP1, protein phosphatase 1; PP2A, protein phosphatase 2A; PP2B, protein phosphatase 2B; SK, small conductance calcium-activated potassium channels; BK, large-conductance calcium-activated potassium channels; PKCδ, protein kinase C delta type; DAG, diacylglycerol; IP: Intrinsic plasticity; LTP: Long term potentiation.

## Materials and Methods

### Animals

Experimental protocols were approved by the Animal Care and Use Committee (A2019-187A, A2018-148A, A2017-060C4) and Gene Recombination Experiment Safety Committee (G2019-020A, 2017-040A) of Tokyo Medical and Dental University. Hetorozygous mice at postnatal day 16–24 (P16–24) of the Aldoc-Venus knock-in line (AldocV, MGI:5620954, Fujita et al., 2014) and the Tg(Rgs8-EGFP)CB132Gsat:C57BL/6N line (Rgs8-EGFP, MGI:3844513, http://www.informatics.jax.org/allele/MGI:3844513) were used. Mice were bred and reared in a specific pathogen-free and 12-12 hour light-dark cycled condition with freely available food and water in the animal facility of the university. Heterozygous Aldoc-Venus pups were obtained by mating male homozygous Aldoc-Venus mice (Fujita et al., 2014; Sarpong et al., 2018) and wild-type C57BL/6N females (CLEA Japan, Tokyo). Heterozygous Rgs8-EGFP mice were obtained from Jax and maintained by mating with C57BL/6N mice. The Rgs8-EGFP mouse is a transgenic reporter line with the transgene containing the coding sequence of enhanced green fluorescent protein (EGFP) inserted into the mouse genomic bacterial artificial chromosome (BAC) at the transcription initiation codon of the regulator of G-protein signaling 8 (Rgs8) gene. PCR genotyping was performed using tail genomic DNA with two primers (Rgs8-108F, TTTAGGTGAGAGGACGTGAGAG; GFP-R, GCGGTCACGAACTCCAGC), which corresponded to the upstream sequence of the GFP insertion site and the coding sequence of the EGFP cDNA of the RGS8-EGFP transgene, respectively (PCR product size, 788 bp; Kaneko et al., 2018). Homozygous male Rgs8-EGFP mice were phenotyped among siblings of mating between heterozygotes Rgs8-EGFP mice. Heterozygous Rgs8-EGFP pups were obtained by mating homozygous Rgs8-EGFP males and wild-type C57BL/6N females.

### Histological procedures

Heterozygous Aldoc-Venus and Rgs8-EGFP pups at P20 were anesthetized with an intramuscular injection of pentobarbital sodium (0.1 mg/g body weight) and xylazine (0.005 mg/g body weight) and perfused transcardially with phosphate-buffered saline (PBS, pH 7.4) with heparin sulfate (0.1%), and then with 4% paraformaldehyde. The brains were dissected in chilled paraformaldehyde and kept in 4% paraformaldehyde for post-fixation and then soaked in sucrose solution (30% with 0.05M phosphate buffer, pH 7.4) for two days. Brains were then coated with gelatin solution (10% gelatin, 10% sucrose in 10mM phosphate buffer, pH 7.4, 32 °C). The gelatin block was hardened by chilling and then soaked overnight in fixative with a high sucrose content (4% paraformaldehyde, 30% sucrose in 0.05 M phosphate buffer, pH 7.4). Serial sections were cut coronally and sagittally using freezing microtome at a thickness of 50 μm. Some sections were mounted on the glass slides, while other sections were rendered to immunostaining for aldolase C. After washing with PBS, floating sections were incubated with the rabbit polyclonal primary rabbit antibody against aldolase C (1:10000, #69076 RRID:AB_2313920, Sugihara and Shinoda, 2004) for two days. After washing with PBS, they were incubated with the donkey secondary antibody against rabbit IgG conjugated with Aleda Fluor 594 (1:400, Jackson ImmunoResearch, 711-585-152) for 1 day. Sections were then mounted on glass slides. Mounted sections were dried and coverslipped with water-soluble mounting medium (CC mount, Sigma C9368-30ML).

Fluorescence images were digitized using a cooled color CCD camera (AxioCam ICm1, Zeiss, Oberkochen, Germany) attached to a fluorescent microscope (AxioImager.Z2, Zeiss) in 12-bit gray-scale with an appropriate filter set. To digitize a section of the cerebellum, 2.5x objective and tiling function of the software to control digitizing (Zen 2 Pro, Zeiss) was used. Images of all serial sections of a brain were obtained with the same exposure parameters. Micrographs were adjusted concerning contrast and brightness and assembled using a software (ZEN 2 Pro, Zeiss and Photoshop 7, Adobe, San Jose, CA, USA). An appropriate combination of pseudo-color was applied to fluorescent images. Photographs were assembled using Photoshop and Illustrator software (Adobe). The software was used to adjust contrast and brightness, but no other digital enhancements were applied.

### Slice preparation

Animals were anesthetized with an intraperitoneal injection of an overdose of pentobarbital (0.1 mg/g, Abbott lab, Chicago, U.S.A.) and xylazine (0.005 mg/g), and euthanized by cervical dislocation. The cerebellar block was dissected from the extracted brain under ice-cold sucrose cutting solution containing (in mM): 87 NaCl, 2.5 KCl, 0.5 CaCl_2_, 7 MgCl_2_, 1.25 NaH_2_PO_4_•2H_2_O, 10 D-glucose, 25 NaHCO_3_, 75 Sucrose and saturated with 95% O_2_ and 5% CO_2_. 200-300 μm parasagittal slices were cut using a vibratome (PRO7, Dosaka, Osaka, Japan). Slices were initially allowed to recover in artificial cerebrospinal fluid (ACSF) solution containing (in mM): 128 NaCl, 2.5 KCl, 2 CaCl_2_, 1 MgCl_2_, 1.2 NaH_2_PO_4_, 26 NaHCO_3_, and 11 Glucose and saturated with 95% O_2_ and 5% CO_2_ at 34°C for 30 min, and then allowed to recover in ACSF at room temperature for at least 1 hour.

### Zebrin stripe identification in slices

In sagittal sections, which were used in experiments of the present study, it was not straightforward to identify zebrin zones. Zebrin zones are not completely parallel to the midsagittal plane, but are tilted laterally in the dorsal position and shifted laterally in the central part of the cerebellum. Furthermore, individual zebrin zones have different widths. For example, zone 1+ ran nearly in parallel with the plane of the slice throughout lobule IV-V in the slice at the midsagittal section. Zones 2+ ran more laterally in the caudal part and more medially in the rostral part of lobule IV-V in the section approximately 400–500 μm from 1+. When cutting sagittal slices, we kept them in order so that the mediolateral position of each slice could be tracked. Because of these morphological properties, which are roughly shown in the unfolded scheme (Sarpong., et al., 2018), zebrin stripes appeared in a characteristic pattern in sagittal sections. Therefore, we could record from Z+ and Z− PCs in identified zebrin zones (1+, 1-, 2+ in lobule IV-V of Aldoc-Venus mice, 5+ and 5- in crus II of Rgs8-EGFP mice).

### Patch-clamp recordings

Slices were placed in the bottom of the recording chamber, soaked in ACSF, and mounted on the stage of a microscope (BX51IW, Olympus, Tokyo, Japan). Slices were then examined with a 10x objective and epifluorescence optics with a filter for appropriate wavelength selection, the distance of the sliced plane from the midline, and location and tilt of Z+ zones were carefully checked. PCs were identified by their characteristic morphology. Z+ zones were visible even in sagittal slices since AldocV mice show a strong contrast of fluorescence expression between Z+ and Z− PCs (Nguyen-Minh et al., 2019). Stripes were then identified regarding the aldolase C stripe pattern which has been mapped upon the unfolded scheme of the cerebellum (Sarpong et al., 2018). The optics of the microscope were changed from epifluorescence to near-infrared Nomarski interference contrast system to approach the PC with the electrode.

Slices were constantly superfused with ACSF at room temperature (24 °C). 100 μM picrotoxin (C0375, Tokyo Chemical Industry Co., Tokyo, Japan) was added to the ACSF to block GABAA channels. PCs were visualized for recording using a 40x water-immersion objective. Patch electrodes (3–5 MΩ) were pulled with a Flaming/Brown type micropipette puller (P-97, Sutter Instruments, San Jose, CA, USA) and filled the internal solution consisting of the following (in mM): 124 K-gluconate, 2 KCl, 9 HEPES, 2 MgCl_2_, 2 MgATP, 0.5 NaGTP, 3 L-Ascorbic Acid, pH adjusted to 7.3 with KOH, osmolarity was adjusted to 280–300 mOsm with sucrose. Signals from the patch pipette were recorded with a MultiClamp 700B amplifier (Molecular Devices, San Jose, USA), digitized at 10–20 kHz and filtered at 2–5 kHz with a Digidata 1440A analog-to-digital converter (Molecular Devices).

The stimulation electrode was made from a glass pipette (0.5–1.0 MΩ) filled with ACSF and a pair of stainless wires placed inside and outside of the pipette. It was positioned in the molecular layer to stimulate PFs. In all experiments, the capacitance was compensated, the impedance bridge was balanced, and bias current (<400 pA) was injected to keep the membrane voltage between −65 and −70 mV. Access resistance was <20 MΩ and both access and input resistances were monitored by applying hyperpolarizing voltage steps (−10 mV) in voltage-clamp mode and −300 pA in current-clamp mode at the end of each sweep, and changed by <20% throughout the recording measured in Clampex (Molecular Devices, San Jose, USA).

### Measurements of intrinsic excitability and intrinsic plasticity of PCs

To evaluate PC excitability, a series of 500-ms long square-shaped current steps of intensity ranging from 0 pA to 1000 pA or 700 pA with increments of 50 pA or 100pA was injected at 7-second intervals from the baseline membrane potential of 70 mV under current-clamp mode. The number of action potentials during the 500 ms period and the time between the first and the last spike (larger than 5mV) were measured for each current injection intensity to obtain the current-spike relationship (cf. Fig. 2A-C).

To induce intrinsic plasticity or LTP of intrinsic stimulation (LTP-IE), we gave tetanizing stimulation, composed of 150 times repetition (every 2 s for 5 min) of a run of five suprathreshold square-shaped depolarizing current injections of 400 pA under current-clamp mode (“LTP-IE protocol”, cf. Fig. 4A, Shim et al.; 2018). We measured the current-spike relationship (above) immediately before the LTP-IE protocol (“baseline”) and 10 and 20 min after the LTP-IE protocol (“10 min”, and “20 min”, cf. Fig. 4F-G). In control experiments, in which the current-spike relationship was measured at the same timing without giving the LTP-IE protocol.

### Measurements of subthreshold afterhyperpolarization (AHP) of PCs

The size of the subthreshold AHP was measured in two ways in PCs from the baseline membrane potential of −70 mV under current-clamp mode. In the synaptic protocol, PFs were stimulated with a train of 5 stimuli of 100 Hz interval to elicit in a PC a summed EPSP of 5–12 mV followed by a subthreshold AHP of 0.5–3.0 mV (cf. Fig. 3A). We normalized the peak amplitude of the subthreshold AHP by dividing it with the peak amplitude of the summed EPSP (“AHP/EPSP ratio”, cf. Fig. 3C). In the current injection protocol, we injected a square current of 50 pA for 500 ms, which produce a depolarization of 3–11 mV, to evoke the subthreshold AHP after the cease of the current injection (cf. Fig. 3B) (Schreurs et al., 1998). The amplitude of the subthreshold AHP was measured from the baseline membrane potential (cf. Fig. 3D).

### Measurements of distinct components of afterhyperpolarization (AHP) of PCs

The fast AHP (fAHP) was measured in the AHP that follows the first action potential evoked by the smallest square current injection among 8 steps (0 pA to 700 pA, 100 pA increment) enough to evoke an action potential (usually 100 pA). Its amplitude was measured from the voltage difference between the negative peak (bottom) of the AHP and the threshold level or the deflection point of the membrane potential (Fig 5A).

To measure the medium AHP (mAFP) and slow AHP (sAHP) we calculated AHP after a burst of action potentials were evoked in a PC by a 300 pA square current injection under current-clamp mode. The cease of the current injection then produced an elongated AHP, which peaked in about 100–200 ms and lasted for about 1 sec. The mAHP and sAHP were measured by calculating the voltage difference between baseline and the negative peak of the elongated AHP and that between the baseline and the decaying AHP 500 ms after the negative peak, respectively (Fig 5B).

### Measurements of the LTP of the PF-PC synaptic transmission

The intensity of the PF stimulation was adjusted to produce about 200-300 pA of EPSC. The amplitude of the EPSC in response to the PF stimulation given every 20s was monitored under voltage-clamp mode and recorded with the online statistic of Clampex. Five minutes after the start of recording, the recording was switched to the current-clamp mode and PF stimulation was given at 1 Hz (LTP induction protocol with the low-frequency PF stimulation) for 5 minutes. Then, the recording was switched back to the voltage-clamp mode and monitoring of the EPSC amplitude was continued for 25 minutes (cf. Fig. 6).

### Statistical analysis

All data were acquired with Clampex 10.7 software (Molecular Devices) and analyzed with Clampfit 10.7 and R programming software. Sample size was determined based on statistical power analysis. Exclusion criteria for patch-clamp experiments are stated above. Statistical significance was determined by using the paired Student’s t-test, two-way ANOVA with repeated measure (within-group comparison of paired events), unpaired Student’s t-test, two-way ANOVA with repeated measure, and the Mann-Whitney U test (between-group comparison), when appropriate. All data are shown as mean ± SEM.

## Acknowledgments

The authors thank Dr. Ryosuke Kaneko for providing us PCR and other information about the Rgs8-EGFP mouse, and Mr. Richard Nana Abankwah Owusu Mensah for proofreading the manuscript. This study was supported by Grant-in-Aid for Scientific Research (KAKENHI) from the Japan Society for the Promotion of Science (19K06919 to IS). VN-M and KT-A are recipients of MEXT scholarship for foreign doctor course students.

## Author Contributions

VN-M and IS: study concept and design and critical revision of the manuscript for important intellectual content. VN-M: acquisition of data. VN-M, KT-A and IS: analysis and interpretation of data. VN-M, and IS: drafting of the manuscript.

